# Single-cell differential expression analysis between conditions within nested settings

**DOI:** 10.1101/2024.08.01.606200

**Authors:** Leon Hafner, Gregor Sturm, Markus List

## Abstract

Differential expression analysis provides insights into fundamental biological processes and with the advent of single-cell transcriptomics, gene expression can now be studied at the level of individual cells. Many analyses treat cells as samples and assume statistical independence. As cells are pseudoreplicates, this assumption does not hold, leading to reduced robustness, reproducibility, and an inflated type 1 error rate.

In this study, we investigate various methods for differential expression analysis on single-cell data, conduct extensive benchmarking and give recommendations for method choice. The tested methods include DESeq2, MAST, DREAM, scVI, the Permutation Test and distinct. We additionally adapt Hierarchical Bootstrapping to differential expression analysis on single-cell data and include it in our benchmark.

We found that differential expression analysis methods designed specifically for single-cell data do not offer performance advantages over conventional pseudobulk methods such as DESeq2 when applied to individual data sets. In addition, they mostly require significantly longer run times. For atlas-level analysis, permutation-based methods excel in performance but show poor runtime, suggesting to use DREAM as a compromise between quality and runtime. Overall, our study offers the community a valuable benchmark of methods across diverse scenarios and offers guidelines on method selection.

## Introduction

Differential gene expression analysis is a fundamental task in single-cell and bulk transcriptomics to detect differences between sample groups. While tools for bulk RNA-seq have matured over the last two decades, there is currently no consensus on differential gene expression analysis on single-cell RNA-seq (scRNA-seq) datasets. In the past, simple statistical tests, such as t-tests or Wilcoxon tests, have commonly been applied to compare expression levels between groups of cells. Since cells of the same sample are not independent observations, this violates a fundamental assumption of most statistical tests, known as *pseudoreplication bias*, which can lead to false-positive results [1, 2].

One solution to avoid pseudoreplication bias is to aggregate the single-cell data by sample and perform statistics on the resulting “pseudobulks” [2]. To retain the advantage of scRNA-seq, this can be done for each cell-type separately. Moreover, linear mixed effects (LME)-models explicitly model the biological replicate as a random effect [1, 3, 4]. Alternatively, resampling-based methods like *distinct* [5] have been proposed. Since Hierarchical Bootstrapping has been applied to similar problems in other scientific fields [6, 7], we adapt Hierarchical Bootstrapping to single-cell data and compare it with LME-models and bulk differential gene expression analysis methods using both simulated and real-world data, covering different analysis scenarios. We provide a comprehensive benchmark, including advanced simulation scenarios, such as atlases, highly unbalanced datasets, and varying proportions of differentially expressed genes. In contrast to previous benchmarks, which concentrated on simpler simulated datasets, our study offers deeper insights into the performance and applicability of these methods in complex biological contexts and guidelines for the community (Fig. 1) [1, 4].

**Fig. 1.**
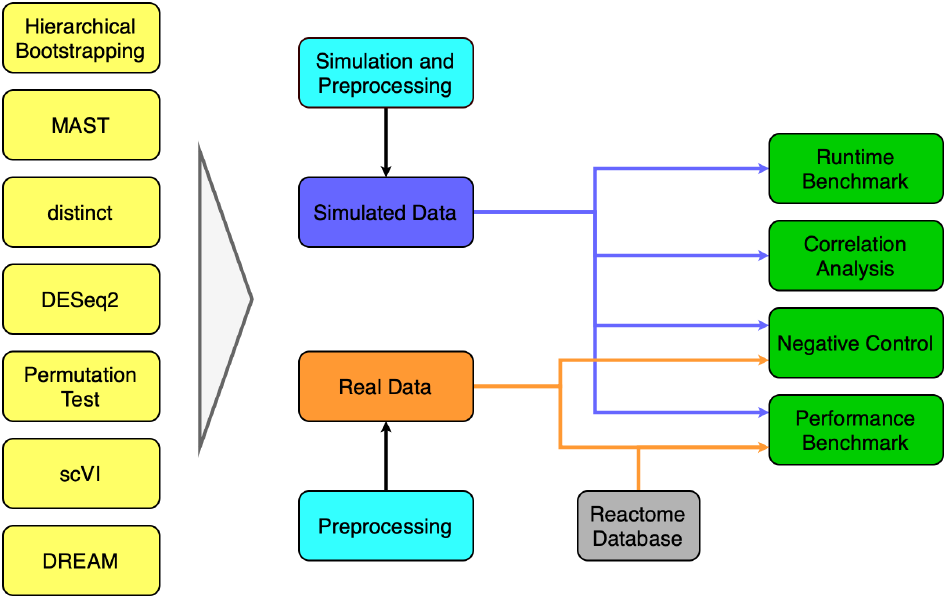
Flowchart of the benchmarking pipeline. Differential gene expression testing methods (yellow) are applied to simulated and real datasets. The data is preprocessed or simulated and preprocessed (light blue). A correlation analysis (supplementary note 1) and a runtime and performance benchmark, as well as a negative control, are then performed (green). For determining the performance on the real dataset, the Reactome database is used (gray) [25].

### Differential gene expression analysis methods

We distinguish between methods that can be applied to single-cell data directly and methods that require pseudobulking to mitigate the pseudoreplication bias [1]. While parametric methods assume a certain data distribution, non-parametric tests can be applied to data independent of its underlying distribution (Tab. 1). If the underlying assumptions are met, parametric tests are considered more powerful than non-parametric ones [8].

*Glossary*

**Linear model** A linear model, or more specifically *linear regression*, estimates the linear relationship between a response variable (*y*) and one or more explanatory variables (*x*_1_, …, *x*_*i*_), by choosing the parameters (*β*_0_, …*β*_*i*_) such that the error term (*ϵ*) in the following equation is minimal: *y* = *β*_0_ +*β*_1_ *x*_1_ +…+*β*_*i*_*x*_*i*_ +*ϵ*. The fundamental assumption is that a constant change in one of the explanatory variables leads to a constant change in the response variable (i.e. “linear relationship”).

**Generalized linear model** A generalized linear model is an extension of linear models that allows for response variables that follow a distribution incompatible with the assumption of a linear relationship, such as binary variables or exponential distributions. Gene expression data is often modeled as a response variable that follows a negative binomial distribution.

**Variance Stabilizing Transform (VST)** For some distributions, the variance depends on the mean. Specifically, in the case of gene expression count data, the higher the gene expression, the higher the variance. VST transforms the data to ensure variance is independent of the mean, allowing the application of simpler statistical models to the data.

**Wald test** The Wald test assesses the statistical significance of a regression coefficient (*β*_*i*_) by comparing the estimated coefficient to its standard error.

**Empirical Bayes shrinkage** Empirical Bayes shrinkage is a technique to improve parameter estimates by leveraging information from all variables (i.e., genes), which is particularly useful when there are not enough replicates to reliably estimate the parameter. An initial estimate of the parameter based on the whole dataset is used as *prior* which is updated using Bayes’ rule with the data seen for the specific variable.

**Random effects** See *mixed effects*.

**Mixed effects** A linear mixed effects model includes both fixed and random effects and is typically used for accounting for nested structures, such as repeated measurements. The fixed effects are captured by explanatory variables’ coefficients like in a standard linear regression model. The random effects part controls for heterogeneity that is not correlated with the explanatory variables.

**Bayes factor** In testing a null hypothesis *H*_0_ against an alternative hypothesis *H*_1_, the null hypothesis is typically rejected if the p-value falls below a significance threshold *α* (typically 0.05). Bayesian statistics, instead, compares the likelihood of two *H*_0_ and *H*_1_. The ratio of the likelihoods is called *Bayes Factor*. Bayes factors can be interpreted more gradually rather than focusing on a single significant threshold, do not require adjustments for multiple hypothesis testing and are arguably more appropriate than p-values for exploratory analyses [13].

### DESeq2

DESeq2 [9] is one of the most widely used methods for differential expression analysis of bulk RNA-seq data. It models gene counts as a negative binomial distribution and applies empirical Bayes shrinkage to calculate gene-wise dispersion estimates – using information from all genes to stabilize the dispersion estimates, especially for genes with a low expression. By default, it uses the Wald test to obtain p-values for each gene. Based on a linear model, it allows complex designs, including continuous and categorical covariates, but no random effects.

### DREAM

DREAM (Differential expression for repeated measures) [10] is an extension of the limma/voom workflow [14] for bulk RNA-seq analysis. On top of continuous and categorical covariates, it allows modeling mixed effects such as multi-experiment data or repeated measures of the same individual. First, this workflow applies a variance-stabilizing transformation (VST) to the gene counts. Then, a linear mixed effects model is fitted to the transformed values, and an empirical Bayes shrinkage is applied to the variance estimates, similar to DESeq2.

### Permutation Test

The Permutation Test is a simple, non-parametric test for comparing the distribution between two groups of samples. It was proposed by Fisher as early as the 1930s [15]. Even though it is agnostic of the underlying distribution, it still assumes independence of the samples. Therefore, it cannot be applied to single cells directly but requires pseudobulk-aggregation. When applied to differential gene expression analysis, the difference in means between the two groups is calculated for each gene. Then, the group labels are randomly shuffled *n* times to obtain a background distribution of the difference in means. The p-value (the probability that an observed difference could have occurred by chance) is the number of permutations that resulted in a greater or equal difference than the one observed divided by *n*. Of note, the smallest p-value achievable by this method is *n*^−1^, making this test computationally expensive, especially in combination with multiple testing correction.

### MAST

MAST [11] is a generalized linear mixed effects model for single-cell data. MAST allows explicitly modeling biological replicates as a random effect, thereby treating single cells as repeated measures of the same sample [1]. It employs a two-component hurdle model that treats single-cell data as “zero-inflated”, i.e. it assumes that zeros are observed more often than expected by a standard distribution. The first component models the discrete expression rate of each gene across cells, and the second component captures the continuous gene expression level conditioned on the expressed gene. MAST expects variance-stabilized counts as input.

### scVI

scVI [12, 16] is an autoencoder for single-cell data, i.e. an artificial neural network that learns a low-dimensional embedding of the single-cell data while removing batch effects and other technical covariates. It has excelled in a benchmark assessing batch-effect removal [17] and has been successfully used to build tissue and disease atlases with *>*1 million cells from heterogenous datasets [18, 19]. From the decoder network, posterior distributions of the corrected gene expression can be retrieved and used for Bayesian differential gene expression analysis. To this end, the probability of two hypotheses (*H*_0_ = “no change” and *H*_1_ = “change”) is calculated and compared using a Bayes factor. The Bayes factor is used to rank the genes instead of a p-value.

### distinct

Distinct [5] is a non-parametric method for differential distribution analysis. Unlike the other methods mentioned in this work that focus solely on changes in the mean, it can detect other changes in the gene expression distribution that do not, or only subtly, change the mean, such as an increase of variance or shifts between bimodal distributions. To do so, it computes an empirical cumulative distribution function (ECDF) for each sample based on the expression of single cells. To compare two groups, the ECDFs of all samples in a group are averaged, and test statistics capturing the distance between the two curves are calculated. A null distribution of the test statistics is obtained using a permutation approach to derive p-values. While distinct can detect changes in distributions beyond their mean, we only use distinct to compare means in this work.

### Hierarchical Bootstrapping

Hierarchical bootstrapping is a non-parametric method for obtaining confidence estimates for differences between groups that has been successfully applied in other fields for dealing with complex, hierarchically structured data [7]. We have adopted this approach to single-cell transcriptomics. Given a dataset with a specified hierarchical structure (e.g. cell < patient < dataset), we randomly sample with replacement for each condition *n* bootstraps with an equal number of cells from each category of all categorical covariates. We obtain empirical gene expression distributions from these samples for both conditions. To obtain a p-value, we count the number of samples in which the first condition exceeds the second condition and vice versa and divide the smaller of the two values by the number of iterations *n*. A more detailed description of our implementation of Hierarchical Bootstrapping, including pseudocode, is available in supplementary note 2. Similar to the Permutation Test, the smallest possible p-value is limited to *n*^−1^. We adopted an adaptive sampling strategy to optimize computational efficiency where, initially, 10 000 bootstrap iterations are drawn for each gene. For genes achieving the smallest possible p-value in this step, the number of iterations is increased to 100 000.

## Benchmark

Since single-cell analysis is a fast-evolving field, we do not include all available methods. Our study covers representative methods across different approaches for single-cell differential expression analysis (i.e. parametric methods like linear models and linear mixed effects models as well as non-parametric methods; see Table 1). Moreover, our benchmark was implemented as a fully reproducible Nextflow pipeline with containerized environments [20], making it easy for method developers to include additional tools.

**Table 1.** Methods partitioned by the statistical framework and used input data.

|  | Parametric | Non-parametric |
| --- | --- | --- |
| Pseudobulk | DESeq2 [9]<br>DREAM [10] | Permutation Test |
| Single-cell | MAST [11]<br>scVI [12] | Hierarchical Bootstrapping<br>*distinct* [5] |

We simulated data using a modified version of the R package splatter with its extension splatPop [21, 22]. To corroborate the findings, we further assess the methods’ performance on experimental data [23, 24], where we consider Reactome pathway genes as ground truth [25]. In addition to the runtime, we assess the false positive rate of the tested tools by simulating data without differentially expressed genes and permute the labels of the experimental data. An overview of all benchmark tasks is provided in figure 1.

## Materials and Methods

The benchmark is implemented as a Nextflow pipeline [20] with predefined containerized environments for each process. Data simulation and plotting, along with the methods DESeq2, distinct, DREAM and MAST are implemented in R. All other methods and processes are implemented in Python. Genes expressed in less than 10% of the cells were removed from the expression matrix because MAST and its linear model require a certain proportion of cells with expression for each gene to work reliably. The individual methods partly require specific preparation steps, where we follow the recommendations in their documentation or vignette. For methods expecting bulk RNA-seq data as input, such as DESeq2, the Permutation Test, and DREAM, we created pseudobulk samples as detailed below.

### Simulation

We selected Splatter and SplatPop as the basis for our simulation due to its extensive customization options, support for nested hierarchies, and comprehensive documentation. We adapted SplatPop with respect to the condition assignment for this work (https://github.com/LeonHafner/sc-guidelines). The simulation process can be divided into two parts. First, we generate the gene means for each sample and the condition-specific expression of the differentially expressed genes. Second, we generate a selected number of cells and expression values for each sample and gene.

#### Dataset scenario

The dataset scenario serves as a balanced baseline with a single batch and consists of two groups (e.g. diseased patient versus healthy control) with 5 samples of 250 cells each. We simulated data sets with 5000 genes of which either 250 (5%) or 25 (0.5%) were differentially expressed between the two conditions. The fold change for the differential expression is drawn from a log-normal distribution with a mean of *μ* = 13.2 and a standard deviation of *σ* = 5.5. The 5% quantile of the values is 6.32 and the 95% quantile is 23.55, thus providing favorable conditions for detecting the vast majority of differentially expressed genes. Unless otherwise noted, these fold-change values apply to this simulation and all other datasets simulated in this work.

#### Atlas scenario

The atlas consists of 3750 cells evenly distributed across three batches of five samples each. Each batch contains at least one sample from each of the two conditions to facilitate comparisons within the batch. The batch effects for the three batches were drawn from log-normal distributions with the following parameters for location and scale: (0.05, 0.1), (0.8, 0.1) and (1.45, 0.05). The resulting log-normal distributions display little to no overlap, ensuring clear differences between batches.

#### Dataset with varying cell numbers scenario

The above scenarios assume a constant number of cells, which does not reflect reality. We hence investigate a modification of the dataset scenario with a variable number of cells per sample taken from a gamma distribution with shape 0.8 and rate 0.0035 and rounded to integer values. These parameters were chosen to ensure that samples with high and low cell counts are simulated, resulting in large differences with a mean cell count of *μ* = 228 and a standard deviation of *σ* = 255. Pseudobulk samples, in particular, typically neglect the number of cells of the contributing samples, possibly leading to an uneven weighting of differential expression. The distribution of the cells per sample is shown in Fig. 2B.

**Fig. 2.**
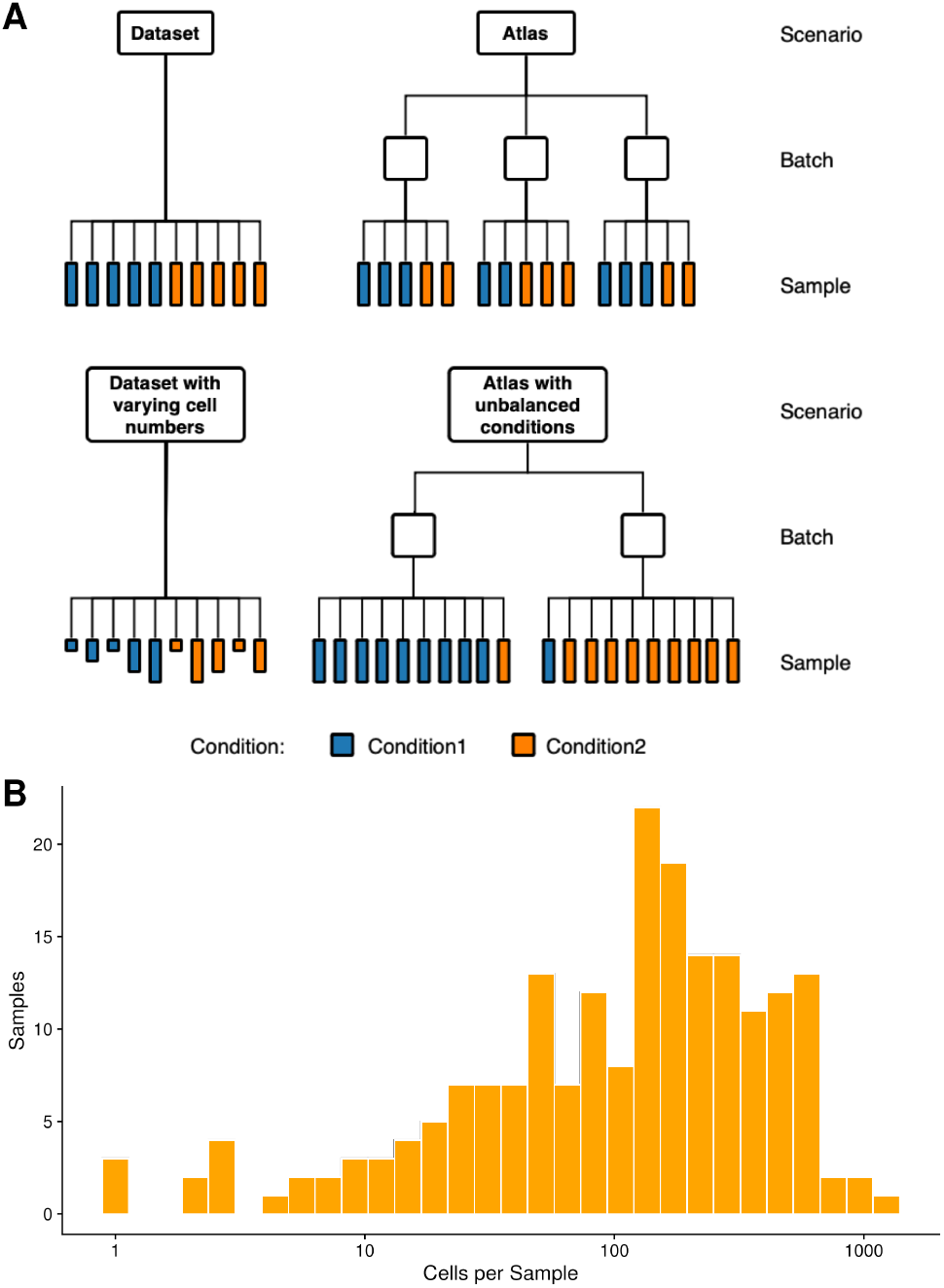
Simulated scenarios. **(A)** shows an overview of the four main scenarios simulated for this study. **(B)** depicts a histogram of the number of cells in all samples of 20 simulations of the dataset with varying cell numbers per sample scenario. These numbers were drawn using a gamma distribution with shape 0.8 and rate 0.0035.

#### Atlas with unbalanced conditions scenario

In the plain atlas scenario, we consider evenly balanced conditions within batches. To assess if methods are biased by strongly unbalanced batches, we modified the Atlas scenario. We considered a batch consisting of nine samples of condition 1 and one of condition 2 and vice versa in a second batch.

### Pathway peturbation dataset

To further assess real-world performance, we used a publicly available single-cell RNA-seq data set with peripheral blood mononuclear cells (PBMC) from 8 patients [24]. Here, each sample was divided into two aliquots before sequencing. One aliquot was stimulated with interferon-beta, while the other served as a control. Since the interferon-beta pathway is well studied, we can use the pathway (obtained from the Reactome database [25], Fig. S1) as a silver standard for differential gene expression analysis. Following the example of the seurat vignette [23], we considered only B cells to avoid cell type bias in the analysis. In addition, genes expressed in less than 10% of cells were removed, resulting in the detection of 1615 genes across 1319 cells.

### Negative Control

To assess how many false-positive genes are detected, we repeated the atlas simulation scenario without conditions, i.e. we expect no significant difference between the arbitrarily assigned conditions. Similarly, we repeated the analysis of the real data set after randomly permuting the condition labels. We count all differentially expressed genes as false positives and assess method performance systematically across different significance thresholds. Since scVI employs a Bayesian statistics instead of generating classical p-values, we evaluated its results at 2000 equidistant cutoffs using the “target FDR” parameter. We relied on scVI’s classification of genes based on these cutoffs.

### Performance Evaluation

Genes we expect to be differentially expressed and recognized by a method are counted as true positives (TP). Similarly, genes that we do not expect to be differentially expressed and are not reported as such are counted as true negatives (TN). Falsely classified genes are counted as false positives or false negatives (FN), respectively [26]. To compare the methods in each scenario, a precision-recall curve (PRC) is used instead of the widely used Receiver-Operating-Characteristic (ROC) because it allows a reasonable interpretation even for very unbalanced data [27, 28]. This is particularly important in this classification problem since only 5% (or 0.5%) of genes are differentially expressed. Moreover, our data may contain genes that show random differential expression. Hence, precision, which focuses on TPs, is likely less biased. We repeated our evaluation of the area of the precision-recall curve (AUPRC) over 20 independent simulations to ensure robustness.

#### Performance Evaluation on real-world data

Since there is no ground truth for the real dataset, and it is not known how strongly each gene is differentially expressed, we rely on the Reactome interferon-beta stimulation pathway as a silver standard (Fig. S1, R-HSA-909733) [25]. 34 of the transcripts from the Reactome database can be found in our dataset. These genes are assumed to be differentially expressed by interferon-beta stimulation and are used as ground truth for creating precision-recall curves.

### Runtime Benchmark

We fixed the number of genes to 1000 and investigated each method’s runtime across an increasing number of cells; vice versa, we fixed the number of cells and increased the number of genes. Each combination was tested 10 times and averaged to obtain the final runtime. We further considered the additional runtime needed for preprocessing and pseudobulking. The runtime was measured using the Unix system time during the execution of each process in the Nextflow pipeline. Each Nextflow task was executed on a single core for comparability, despite some methods supporting multi-core usage. The HPC used for generating the figures consists of several machines, each equipped with an Intel Xeon Gold 6148 processor with a clock speed of 2.40 GHz.

### Method implementations

We have structured the methods as separate processes within a Nextflow pipeline, where each method has its own containerized environment tailored to meet the recommended use.

#### DESeq2

The raw simulation data without normalization was used as input. The design matrix for DESeq2 varied among the different scenarios. For the Atlas scenarios and also on the real data, batch and condition information were included:

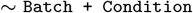

For the Dataset scenarios, only the information about the condition was provided:

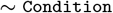

We also assessed the method’s performance on the Dataset with varying cell numbers per sample by including the logarithmized number of cells per pseudobulk (PB) as a fixed effect in the design matrix:

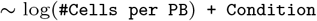

In the first step, a DESeq2 dataset based on the counts was created. Size factors were calculated prior to differential expression analysis. We used R version 4.2.3, DESeq2 1.38.3 and anndataR 0.99.0, to transfer the gene expression data between Python and R.

#### Permutation Test

The Permutation Test was implemented using counts per million (cpm) normalized single-cell count data as input. Our implementation allows users to specify the minimum and maximum number of iterations (10 000 and 100 000 by default) as parameters. We permuted the condition labels of all cells 10 000 times and compared these permutations to the initial condition assignment. If none of these permutations resulted in an assignment more extreme than the initial one, we adaptively increased the number of iterations to 100 000, seeking at least one more extreme permutation. Should no permutation be more extreme than the initial assignment, the method outputs the p-value 0, which indicates a value below 1 *×* 10^−5^. We used Python 3.11.9, Scanpy 1.10.1, Anndata 0.10.7, Pandas 2.2.2 and NumPy 1.26.4 to process the data and implement the Permutation Test.

#### distinct

The data preprocessing adhered to the guidelines provided by the developer of the distinct package. Log-normalized counts and a design matrix were used as inputs for the primary function of distinct. The design matrix included batch and condition assignments for the Atlas and the real scenario; only the condition assignment was used for the Dataset scenarios. The globally adjusted p-values derived from the distinct output were propagated throughout the analysis pipeline. We conducted benchmarks using distinct 1.10.0. To prepare log-normalized counts, we utilized R 4.2.3, scuttle 1.8.4 and SingleCellExperiment 1.20.1. Additionally, anndataR 0.99.0 facilitated the conversion of the data into R objects.

#### DREAM

DREAM (Differential expression for repeated measures) is a method for differential expression testing on bulk RNA-Seq data and is therefore applied to pseudobulked single-cell data [10]. The expression data processing mirrored the approach outlined in the DREAM vignette. A `DGEList` object was created from the raw counts, from which normalizing factors were derived. In the case of the Atlas scenarios, a random effect was modeled for the batches as follows:

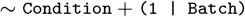

For the dataset scenarios, only the condition was modeled:

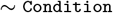

For the Dataset with varying cell numbers per sample we included the logarithmized number of cells per pseudobulk as an additional factor into the design matrix:

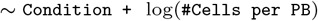

Due to pseudobulking, there was no need to model the sample level separately, as each sample and gene combination yielded a single expression value. The `voomWithDreamWeights` function was used to generate a DREAM object, and the data analysis was conducted using the `dream` and `eBayes` functions. Limma was finally used to extract the differentially expressed genes. The p-value produced in the output table was used for further analyses. For benchmarking, we used R 4.2.3 and the variancePartition package version 1.28.9. The data processing relied on data.table 1.14.8, edgeR 3.40.2 and SingleCellExperiment 1.20.1. AnndataR 0.99.0 was employed to transfer the data from Python to R.

#### MAST

MAST received log-transformed counts as input. For the dataset scenarios, sample information was included and modeled as a random effect. This modeling approach is analogous to pseudobulking and counteracts the pseudoreplication bias. This results in the following model:

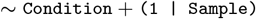

The batch effects of the atlases were modeled as a random effect:

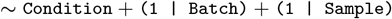

While these could theoretically be modeled as fixed effects, that would require that each batch includes at least one sample from each condition to be valid. After fitting the model, t he hurdle model’s p-value and logarithmic fold changes were extracted for the condition coefficient. R 4.2.3 and MAST 1.24.1 were used. AnndataR 0.99.0 facilitated the conversion of gene expression objects from Python to R.

#### scVI

ScVI was trained directly on the raw single-cell expression data, following the guidelines outlined in the scVI vignette. The `setup anndata` function was used to create an instance of the model. Both Batch and Sample were specified as categorical keys for the atlas scenarios and the real-world dataset. For the dataset scenarios, only the Sample covariate was used. The mini-batch size was dynamically increased if splitting the data resulted in a mini-batch containing only a single gene. The trained model was evaluated by grouping the data according to condition values. The probability of a gene not being differentially expressed was treated as the p-value for further analyses, with the exception of the negative control. For the negative control, following the advice of the scVI developers, we counted the differentially expressed genes at 2000 equidistant cutoffs using scVIs “ target FDR” parameter and relied on its classification of positives and negatives. The method also adjusts those values to account for multiple testing issues. Additionally, as the scVI developers also recommend filtering the data for highly variable genes before analysis, we conducted further benchmarks across all methods on datasets that had been filtered to retain only the top 10% of genes with the highest variability. Genes simulated as differentially expressed that were excluded by this filtering process were assessed as false negatives. For this filtering process, the implementation of Seurat_v3 in Scanpy was employed. The benchmark utilized Python 3.11.9 and scVI 1.1.2. Data preprocessing was performed using pandas 2.2.2 and scanpy 1.10.1.

#### Hierarchical Bootstrapping

We implemented Hierarchical Bootstrapping in Python. The cpm normalized single-cell counts serve as input for the method. Additionally, one can specify the hierarchy of the data, the sample size and the aggregation function. The bootstrapping step and the evaluation step are encapsulated in two separate functions. The first one generates samples according to the specified hierarchies and sample sizes, aggregates them and stores them in a new Anndata object. The evaluation step iterates over this object and counts how often the first condition generated higher expression values than the second condition and vice versa. The gene-wise p-value is obtained by the following formula:

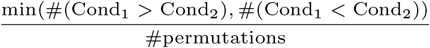

We used Python 3.11.9, Scanpy 1.10.1, Anndata 0.10.7, Pandas 2.2.2 and NumPy 1.26.4 to implement the Hierarchical Bootstrapping method.

## Results

### Performance evaluation on simulated scenarios

Before utilizing the simulated data for benchmarking, we conducted a correlation analysis. This analysis confirmed the presence of pseudoreplication bias in the simulated data, as the transcriptomes of cells within a sample were more highly correlated than those of cells between samples (supplementary note 1). We then evaluated the performance of the methods on the simulated scenarios described in Fig. 2A. Most methods performed well on the dataset scenario, with DESeq2 leading the field with a mean AUPRC of 0.93 followed by scVI with a mean AUPRC of 0.87 and the remaining methods ranging between a mean AUPRC of 0.71 and 0.80 (Fig. 3A).

**Fig. 3.**
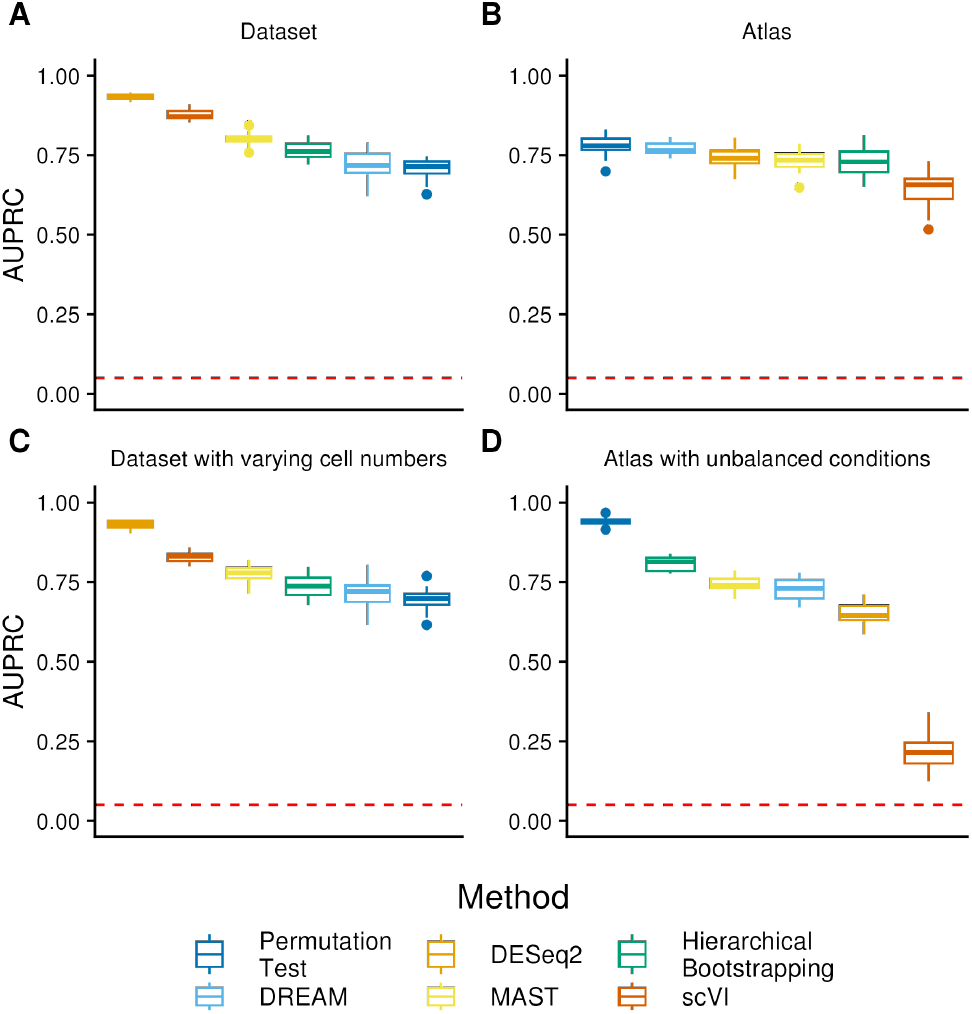
Performance of the methods on the simulated data scenarios. Performance of the methods on the simulated scenarios over 20 independent simulations and runs each for **(A)** the Dataset scenario, **(B)** the Atlas scenario, **(C)** the Dataset scenario with varying cell numbers, and **(D)** the Atlas scenario with unbalanced conditions. The methods were sorted based on their performance. The random baseline is indicated by the dashed line. Full precision-recall curves for a randomly selected run are shown in Figure S7. We excluded distinct from those benchmarks, as it computed the lowest possible p-value for the majority of the genes and therefore did not provide useful results.

**Fig. 4.**
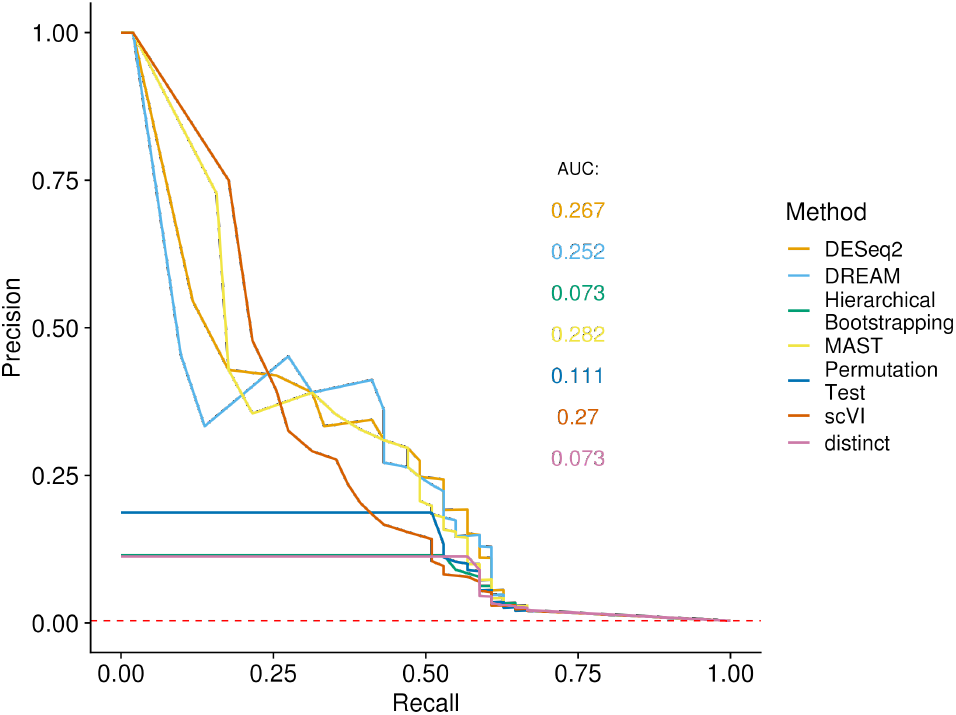
Precision-recall plot of the performance on a real data set. Performance of all methods on a single-cell dataset of 1319 B-cells stimulated with interferon beta. The corresponding Reactome pathway serves as the ground truth for determining the precision-recall curve.

In the Atlas scenario, the methods generally performed worse, likely owing to the batch effects. Here, the Permutation Test performs best with an AUPRC of 0.78. DREAM, DESeq2, Hierarchical Bootstrapping and MAST perform comparably with a mean AUPRC between 0.73 and 0.77 while scVI showed a significantly worse performance AUPRC of 0.66 (Fig. 3B). ScVI’s poor performance in this scenario is surprising, as the batch information was provided using the batch key parameter and sample information was passed using the categorical covariates parameter.

Since making pseudobulks from samples with identical cell numbers seems unrealistic in practice, we used the Dataset with varying cell numbers scenario to ask how different cell counts per sample impact the results. Overall, the results are comparable with the regular dataset scenario, except the performance of scVI, which is slightly worse (Fig. 3C). Therefore, we conclude that at least within the range of cells per sample tested here, varying numbers of cells per sample only have a minor impact on the results of a pseudobulk DE workflow. For the parametric pseudobulk methods, we additionally tried passing the logarithmized number of cells in a pseudobulk as a continuous, fixed effect covariate to the methods. However, this resulted in significantly worse performance (Fig. S2).

It is not always possible to obtain single-cell data in a well-controlled experimental setting with (almost) equal number of samples in each condition, especially when compiling a single-cell atlas from heterogeneous data sources [19]. In the atlas with unbalanced conditions scenario, we tested how methods perform in an extreme case, with only one sample per batch being from the first or second condition, respectively. As in the atlas scenario, the Permutation Test achieves the best result with a mean AUPRC of 0.94, followed by Hierarchical Bootstrapping with an AUPRC of 0.81. MAST and DREAM achieve a mean AUPRC between 0.73 and 0.74, DESeq2 of 0.65 and scVI performs the worst with an AUPRC of 0.22 (Fig. 3D). While some methods show a better performance on the unbalanced atlas than on the balanced atlas, this might be due to the different number of batches. Noticeably, the two non-parametric methods (Permutation Test and Hierarchical Bootstrapping) have an edge over parametric methods in this scenario, while in particular, scVI appears to have difficulties. The reason might be that in this extreme case (only one sample per batch for one of the groups), parametric methods struggle to obtain reliable parameter estimates. While the Permutation test doesn’t have these issues, its good performance was still surprising to us, as it doesn’t account for batch effects, which could, in theory, lead to false-positive results when the change in gene expression is contradicting between the two datasets (e.g. downregulated in batch 1 and upregulated in batch 2). An inspection of the simulated gene expression revealed that very little such contradicting cases are present within the set of the truly differentially expressed genes (Fig. S3), therefore, explaining the good performance of the simplistic Permutation Test.

We additionally examined the change in performance with only 0.5% (instead of 5%) differentially expressed genes (Fig. S4). Expectedly, the performance of all methods decreased. In the dataset scenario, parametric method tended to be less affected than the non-parametric ones. Furthermore, we determined the performance of the methods on filtered datasets using only 10% of the genes with the highest variability, which was recommended by the authors of scVI. However, this led to a decreased performance of all methods, including scVI (Fig. S5).

Distinct’s performance didn’t outperform random baseline in any of the scenarios (Fig. S6). We observed that running distinct resulted in a p-value of 0.0001 for 63% to 89% of the genes, which is the minimal achievable p-value at default setting limited by the number of permutations.

### Evaluation on pathway perturbation dataset

We applied the methods to a single-cell dataset of 1319 B-cells partially stimulated with interferon-beta, considering the genes in the corresponding Reactome pathway as differentially expressed. In this setting, MAST performs best with an AUPRC of 0.282, closely followed by scVI, DESEq2, and DREAM with an AUPRC between 0.25 and 0.27. The Permutation Test achieves an AUPRC of 0.11, while both Hierarchical Bootstrapping and distinct achieve an AUPRC of 0.073. Notably all three non-parametric methods perform worse than the parametric methods on this scenario. We acknowledge the limitations of using an imperfect ground-truth. Reactome contains all genes known to be involved in interferon-beta stimulation. While this pathway is well-studied, there might still be further genes involved. Additionally, secondary genes outside the pathway could be perturbed, too. Moreover, not all genes necessarily alter their expression level upon pathway stimulation, or could be regulated at protein rather than mRNA level. In this light it is not surprising that the methods achieve a significantly worse performance than in a well-controlled simulation experiment. Nevertheless, it remains informative to assess the relative ranking in this setting.

### Negative control shows that not all methods are well-calibrated

In this section, we performed a “negative control” benchmark to assess if the methods appropriately control for false discoveries when no differentially expressed genes are present by comparing p-value cutoffs to the observed false discovery rate. This allows us to detect whether the methods were overly conservative in their classification. DREAM and the Permutation test aligned very well with the diagonal in the simulated data (Fig. 5), making them neither too conservative nor too liberal. DESeq2 and MAST slightly exceed the nominal FDR in the range of small p-values. scVI is over-conservative for p-values < 0.03 but exceeds the nominal FDR for all cutoffs above. Hierarchical bootstrapping exceeds the nominal FDR at all cutoffs. For the real dataset with permuted labels (Figure S8B), the trends were similar; the Permutation Test and DESeq2 appeared slightly conservative, while DREAM and MAST controlled their FDR reasonably well.

**Fig. 5.**
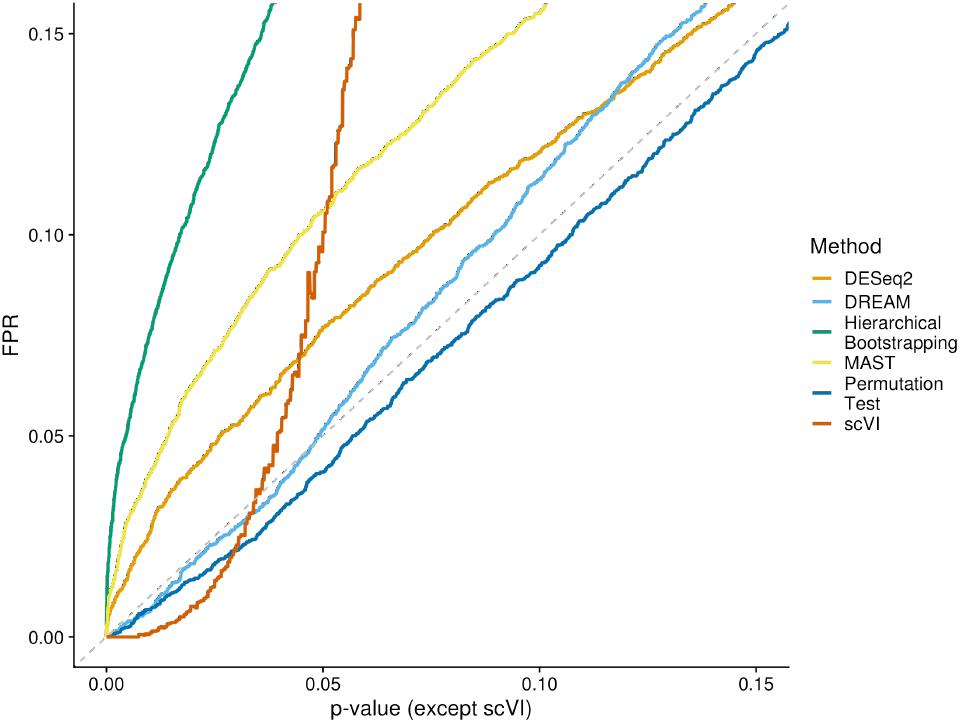
Negative control based on raw p-value cutoff and false positive rate (FPR). This plot shows p-value cutoffs versus the observed false positive rate (FPR, defined as the proportion of genes falsely classified as positive). In a perfect case, the observed FDR corresponds to the p-value cutoff. Values below the diagonal indicates that a method is over-conservative, while values above the diagonal indicate that a method doesn’t appropriately control for false discoveries. Since scVI does not generate classical p-values but employes Bayesian statistics, we passed 2000 equidistant cutoffs to scVI via its “target FDR” parameter and relied on scVI’s classification of positive and negatives. This plot only shows the results up to a p-value of 0.15, the plot over the entire axis range can be seen in Fig. S8A.

### Runtime benchmark

We first varied the number of genes, while maintaining a constant cell count of 1000 (Fig. 6A). DESeq2 and DREAM display only minimal linear increase in runtime, despite a hundredfold increase in the number of genes. In contrast, all other methods exhibit a steeper, yet still linear, increase in runtime with higher gene numbers. The Permutation Test is slowest taking almost 5 hours to run for 10 000 genes, followed by Hierarchical Bootstrapping, MAST, distinct and scVI.

**Fig. 6.**
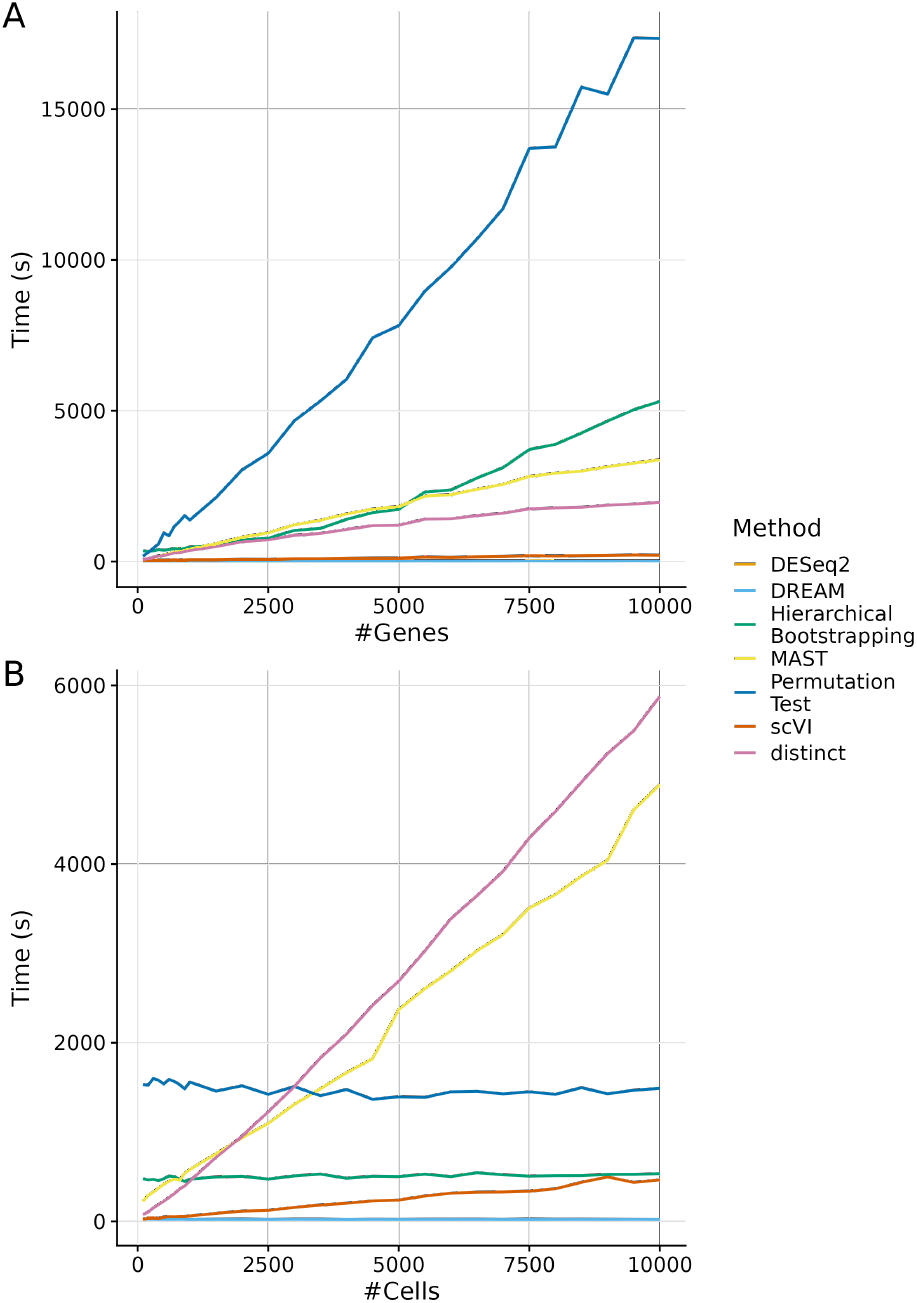
Runtime comparison of the methods. The plots show the runtime of the different methods averaged over ten independent runs. Although pseudobulking had a minimal impact on the overall runtime, it was included in the assessments for DESeq2, the Permutation Test, and DREAM, as these methods work with pseudobulked data. **(A)** Runtime on data simulated to have a fixed cell count of 1000 and between 100 and 10 000 genes. **(B)** Runtime on data simulated to have a fixed gene count of 1000 and between 100 and 10 000 cells.

**Fig. 7.**
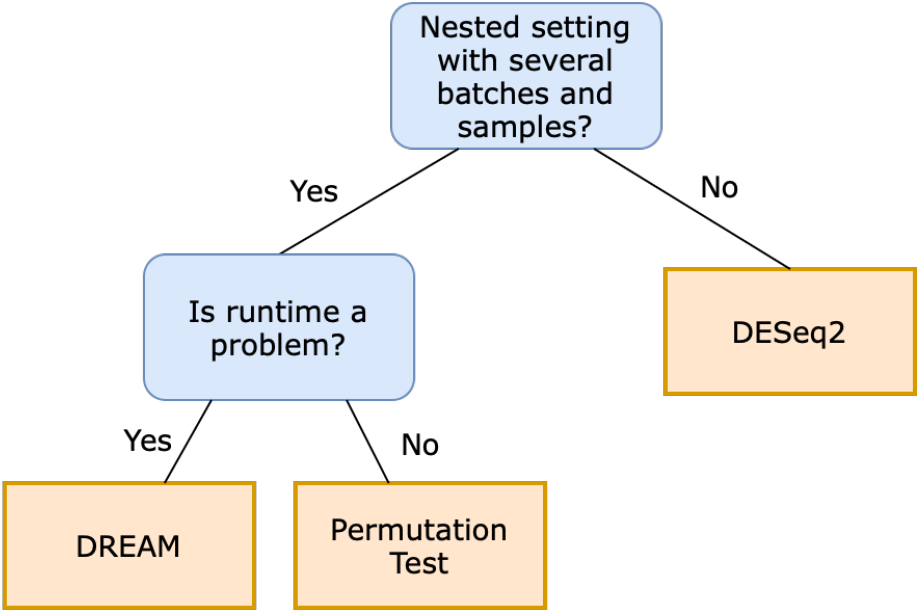
Decision tree supporting users in selecting an appropriate method for single-cell differential expression analysis.

Next, we evaluated the performance with a constant number of genes and an increasing number of cells (Fig. 6B). DESeq2, DREAM, the Permutation Test and Hierarchical Bootstrapping demonstrated constant runtimes, while distinct, MAST and scVI showed a linear increase in runtime depending on the number of cells. DESeq2, DREAM and the Permutation test are independent of the number of cells, because they work with pseudobulked data. Hierarchical bootstrapping samples a fixed number of bootstrapping iterations from the cell pool, making it largely independent of the pool size. Its runtime would likely increase with additional bootstrapping iterations and samples per hierarchical level. MAST, distinct and scVI work on single-cells and therefore depend on the cell count, however scVI is significantly faster than the latter two methods.

Overall, as expected, resampling-based methods were slower than parametric methods and pseudobulk-methods were faster than single-cell methods, with the exception of scVI which is almost as fast as DESeq2 and DREAM, despite working on single cells.

## Discussion and Conclusion

This work explores the performance of differential gene expression analysis methods in hierarchically structured single-cell datasets such as atlases. A novel contribution is the adaptation of the Hierarchical Bootstrapping method, previously successful in neuroscience, to single-cell biology. This method and six other current methods were thoroughly investigated for its performance in the same scenarios. The aim was to provide recommendations for the best method.

Four simulated scenarios and a real dataset were used for benchmarking and their effects on the methods were investigated.

While this work thoroughly examines the most common scenarios in single-cell analysis, it’s important to acknowledge its limitations. For instance, it could be further explored whether the performance of the methods decreases continuously as the number of batches increases or whether methods that have not shown the best performance so far work better with larger batch numbers. Another limitation is that, in reality, single-cell datasets typically encompass a substantially greater number of cells than what we have simulated here. Certain benchmarked methods have already reached the upper limits of their scalability, making it challenging to increase the simulated cell number. In addition, further research could be done regarding the cell number of a pseudobulk. Knowing the minimum reasonable number of cells in a pseudobulk would be particularly interesting. One must also be aware of the uncertainty inherent in using simulated data. Some methods might excel due to their concordance with simulated data characteristics. For instance, drawing expression values from distributions may favor parametric methods. We also note that methods such as MAST assume zero-inflated counts which do not represent single-cell data as well as previously thought [29]. Regarding the real dataset, additional evaluation methods would also be possible. For example, genes considered to be differentially expressed could also be found in the literature or data sources other than the Reactome database could be included. Nevertheless, when comparing differential expression methods, the main focus will be on simulated data, since only for these the real differential expression is known and real biological data is rather laborious to generate or may contain too many confounding factors to perform a meaningful benchmark.

Based on our comprehensive research, we provide a practical guideline to the community. This guideline is specifically designed to empower researchers, scientists, and practitioners in the field of single-cell biology and gene expression analysis to select the most appropriate method for their specific dataset and research objectives. First, a user should consider whether their data contains a nested hierarchy or whether it is a planar dataset with individual samples. If the latter is the case, the current state-of-the-art method DESeq2 should be used after pseudobulking, as it delivers the best results on the dataset scenario and performs well on the real-world dataset combined with a very fast runtime. If the data consists of a nested hierarchy, the user should consider the number of genes as this massively impacts runtime. For increasingly larger data sets, the user should consider DREAM. If the data has a small number of genes, then the Permutation Test can be used for both balanced and unbalanced batches.

In conclusion, pseudobulking, followed by classical methods for bulk RNA-seq data, is often the most effective strategy. More sophisticated methods tailored toward single-cell data can add benefits in complex scenarios such as the integration of multiple unbalanced batches. Since atlas-level data integration becomes more prevalent, further development in refining such methods is warranted. Some methods, such as MAST, can not cope with current atlas-level data sets with millions of cells. Our extensible Nextflow benchmarking framework allows researchers to quickly evaluate new methods with respect to the state-of-the-art, thus facilitating rapid and continuous method evaluation in a rapidly moving field.

## Data availability

All figures and results presented in this paper can be reproduced using our Nextflow pipeline with containerized environments, which is available on GitHub: https://github.com/LeonHafner/sc-guidelines Figures and Singularity containers are additionally available for download from Zenodo: https://doi.org/10.5281/zenodo.12544494

## Acknowledgements

We thank Nir Yosef and his lab for their valuable review and feedback on our use of scVI in this study.

