## Supplement for "Single-cell differential expression analysis between conditions within nested settings"

**Supplementary Note 1: Correlation Analysis**

Consistent with the notion of pseudoreplication bias, transcriptome profiles of cells within a sample show higher pairwise correlation compared to two cells from different samples [1]. To confirm that our simulated data showed this behavior, we simulated 10 samples with 250 cells each in a single batch. In this analysis, we first excluded redundant genes with more than 0.1 absolute Spearman correlation and then randomly sampled 500 genes for further analysis. One cell was drawn from each sample, and then the pairwise correlations of the drawn cells were computed to calculate the between-sample correlations. This procedure was repeated 1,000 times, resulting in a total of 45,000 between-sample correlations. For the within-sample correlations, we sampled 4,500 cell pairs (without replacement) within each sample, and computed their Spearman correlations to obtain a total of 45,000 within-sample correlations. This resulted in the within- and between-sample correlations shown in Fig. S9.

The correlation of cells within a sample for each dataset is higher than the correlation across samples. Thus, it can be concluded that cells in the simulated data are not statistically independent and would introduce pseudoreplication bias if not accounted for. This is consistent with the evidence found for real datasets

**Supplementary Note 2. Pseudocode for the Hierarchical Bootstrapping method on single-cell data.**

The first part of the pseudocode describes the bootstrapping step, where we cycle through the number of bootstrapping iterations and draw sample cells based on the specified hierarchy. The second part is called the evaluation step and iterates over the data aggregated in the bootstrapping step. It generated the p-value of a gene by counting in which of the conditions it has a higher expression more frequently.

```
// First step: Bootstrapping phase
output = {}
for i = 1 to n do
    for each condition do
        resample first hierarchy level with sample size = |{hierarchy level 1}|
        restrict the selection to the drawn samples
        for each of the remaining hierarchy levels do
            resample hierarchy level with user specified sample size
            restrict the selection to the drawn samples
        end
        aggregate selected samples with specified aggregation function
        add results to output
    end
end

// Second step: Evaluation phase
data := output
pvalue = {}
for each gene do
    condition1_higher = 0
    condition2_higher = 0
    for i = 1 to n do
        if data[i, condition1] > data[i, condition2] then
            condition1_higher += 1
        else
            condition2_higher += 1
        end
    end
    pvalue[gene] = min(condition1_higher / n, condition2_higher / n)
end
```

*Table S1. Results of runtime benchmarking with varying number of genes. The runtimes (in seconds) of the methods MAST, distinct, DESeq2, Permutation Test, Hierarchical Bootstrapping (hb), scVI, and DREAM are shown. In addition, the runtime for simulation and pseudobulking (pb) is depicted. For the runtime graphs, the time required for pseudobulking was added to the runtime of DESeq2, the Permutation Test and DREAM since these require pseudobulked data.*

| **n_genes** | **n_cells** | **sim** | **pb** | **mast** | **distinct** | **deseq2** | **permutation** | **hb** | **scvi** | **dream** |
|---|---|---|---|---|---|---|---|---|---|---|
| 100 | 1000 | 29.94 | 4.41 | 54.53 | 74.65 | 20.10 | 174.46 | 358.76 | 43.15 | 18.67 |
| 200 | 1000 | 29.02 | 4.16 | 93.25 | 107.16 | 18.92 | 296.03 | 342.14 | 44.16 | 17.36 |
| 300 | 1000 | 28.59 | 4.10 | 134.14 | 144.15 | 18.98 | 437.50 | 357.94 | 43.93 | 16.46 |
| 400 | 1000 | 30.02 | 4.31 | 184.94 | 189.07 | 20.15 | 580.39 | 396.54 | 53.45 | 17.36 |
| 500 | 1000 | 30.51 | 4.22 | 217.28 | 207.65 | 18.82 | 948.75 | 377.58 | 51.74 | 16.80 |
| 600 | 1000 | 30.50 | 4.29 | 268.28 | 242.31 | 19.36 | 854.74 | 399.23 | 54.15 | 16.37 |
| 700 | 1000 | 31.23 | 4.47 | 309.87 | 289.04 | 20.08 | 1156.48 | 432.90 | 62.00 | 17.08 |
| 800 | 1000 | 31.05 | 4.26 | 337.23 | 296.56 | 19.28 | 1325.13 | 417.50 | 53.93 | 16.26 |
| 900 | 1000 | 30.92 | 4.12 | 381.33 | 331.39 | 19.01 | 1521.79 | 434.53 | 56.34 | 16.10 |
| 1000 | 1000 | 32.84 | 4.59 | 425.66 | 372.78 | 21.01 | 1373.66 | 487.07 | 63.90 | 16.99 |
| 1500 | 1000 | 32.27 | 4.10 | 594.35 | 496.48 | 19.09 | 2117.92 | 532.59 | 64.05 | 16.16 |
| 2000 | 1000 | 35.50 | 4.49 | 812.80 | 662.02 | 20.95 | 3035.82 | 726.20 | 78.41 | 17.55 |
| 2500 | 1000 | 33.81 | 4.08 | 953.51 | 721.92 | 19.88 | 3585.70 | 777.25 | 75.60 | 16.01 |
| 3000 | 1000 | 37.24 | 4.47 | 1218.05 | 870.54 | 21.27 | 4660.22 | 1033.40 | 96.77 | 17.75 |
| 3500 | 1000 | 35.60 | 4.10 | 1372.22 | 933.59 | 19.94 | 5320.79 | 1100.95 | 97.82 | 16.29 |
| 4000 | 1000 | 36.93 | 4.17 | 1578.51 | 1066.74 | 20.84 | 6037.05 | 1401.18 | 112.92 | 17.00 |
| 4500 | 1000 | 39.33 | 4.32 | 1740.50 | 1193.55 | 21.07 | 7415.66 | 1622.54 | 121.20 | 17.61 |
| 5000 | 1000 | 36.78 | 4.25 | 1828.64 | 1206.80 | 20.70 | 7816.86 | 1733.96 | 122.72 | 16.83 |
| 5500 | 1000 | 40.87 | 4.64 | 2168.41 | 1408.51 | 22.43 | 8958.25 | 2306.74 | 151.84 | 18.48 |
| 6000 | 1000 | 39.53 | 4.19 | 2218.31 | 1419.10 | 21.46 | 9754.56 | 2376.30 | 141.27 | 17.05 |
| 6500 | 1000 | 41.72 | 4.38 | 2396.07 | 1524.01 | 22.27 | 10693.01 | 2771.10 | 157.94 | 17.28 |
| 7000 | 1000 | 40.74 | 4.24 | 2561.71 | 1603.12 | 21.55 | 11698.24 | 3116.40 | 169.71 | 17.90 |
| 7500 | 1000 | 44.13 | 4.61 | 2818.79 | 1749.01 | 24.28 | 13691.40 | 3716.01 | 197.50 | 18.78 |
| 8000 | 1000 | 43.31 | 4.49 | 2929.63 | 1780.76 | 23.80 | 13736.05 | 3886.08 | 188.50 | 17.82 |
| 8500 | 1000 | 45.04 | 4.23 | 2998.42 | 1797.89 | 23.44 | 15725.48 | 4263.53 | 200.24 | 17.91 |
| 9000 | 1000 | 44.76 | 4.29 | 3145.61 | 1871.13 | 23.17 | 15489.87 | 4658.51 | 208.02 | 17.81 |
| 9500 | 1000 | 43.50 | 4.01 | 3261.59 | 1909.15 | 23.14 | 17352.88 | 5035.23 | 220.77 | 20.64 |
| 10000 | 1000 | 44.43 | 3.97 | 3376.53 | 1960.28 | 23.39 | 17333.21 | 5309.40 | 213.99 | 17.54 |

*Table S2. Results of runtime benchmarking with varying number of cells. The runtimes (in seconds) of the methods MAST, distinct, DESeq2, Permutation Test, Hierarchical Bootstrapping (hb), scVI, and DREAM are shown. In addition, the runtime for simulation and pseudobulking (pb) is depicted. For the runtime graphs, the time required for pseudobulking was added to the runtime of DESeq2, the Permutation Test and DREAM since these require pseudobulked data.*

| **n_genes** | **n_cells** | **sim** | **pb** | **mast** | **distinct** | **deseq2** | **permutation** | **hb** | **scvi** | **dream** |
|---|---|---|---|---|---|---|---|---|---|---|
| 1000 | 100 | 36.91 | 6.86 | 222.38 | 75.43 | 21.05 | 1525.63 | 478.77 | 21.07 | 17.94 |
| 1000 | 200 | 39.02 | 6.53 | 290.65 | 103.28 | 21.00 | 1517.41 | 464.82 | 37.81 | 20.94 |
| 1000 | 300 | 39.11 | 5.27 | 327.33 | 149.41 | 21.16 | 1594.45 | 470.72 | 39.48 | 20.91 |
| 1000 | 400 | 34.96 | 5.39 | 371.92 | 188.67 | 20.35 | 1574.20 | 456.12 | 34.86 | 18.39 |
| 1000 | 500 | 36.51 | 4.67 | 416.09 | 225.08 | 21.95 | 1532.28 | 478.04 | 48.45 | 18.23 |
| 1000 | 600 | 37.65 | 4.72 | 448.13 | 267.46 | 22.66 | 1584.39 | 507.69 | 53.42 | 18.42 |
| 1000 | 700 | 35.33 | 4.61 | 473.68 | 309.27 | 21.36 | 1562.03 | 501.47 | 51.77 | 18.75 |
| 1000 | 800 | 33.30 | 4.43 | 461.20 | 359.04 | 20.45 | 1527.83 | 471.59 | 55.68 | 21.58 |
| 1000 | 900 | 33.82 | 4.49 | 537.26 | 396.85 | 20.28 | 1478.62 | 447.07 | 56.50 | 18.50 |
| 1000 | 1000 | 32.12 | 4.23 | 579.81 | 451.57 | 20.02 | 1554.52 | 472.71 | 61.28 | 16.99 |
| 1000 | 1500 | 35.67 | 4.83 | 756.94 | 716.51 | 21.50 | 1453.86 | 497.41 | 88.81 | 17.87 |
| 1000 | 2000 | 35.99 | 4.60 | 935.18 | 955.41 | 23.16 | 1513.09 | 505.07 | 114.69 | 17.61 |
| 1000 | 2500 | 36.00 | 4.18 | 1095.89 | 1225.99 | 20.32 | 1418.64 | 473.81 | 122.86 | 17.39 |
| 1000 | 3000 | 37.21 | 4.52 | 1308.67 | 1508.45 | 22.51 | 1505.15 | 509.36 | 156.23 | 20.11 |
| 1000 | 3500 | 38.67 | 4.90 | 1483.86 | 1829.46 | 22.24 | 1401.60 | 529.49 | 182.85 | 17.67 |
| 1000 | 4000 | 37.77 | 4.22 | 1663.13 | 2096.18 | 19.75 | 1472.58 | 482.78 | 203.06 | 16.85 |
| 1000 | 4500 | 41.22 | 4.95 | 1820.08 | 2419.59 | 20.83 | 1361.44 | 505.90 | 229.47 | 18.05 |
| 1000 | 5000 | 39.81 | 4.37 | 2370.21 | 2688.93 | 20.53 | 1390.98 | 502.21 | 238.62 | 17.91 |
| 1000 | 5500 | 41.70 | 4.87 | 2604.21 | 3027.27 | 21.45 | 1383.86 | 528.20 | 283.38 | 18.57 |
| 1000 | 6000 | 42.23 | 4.67 | 2803.79 | 3388.38 | 21.67 | 1446.13 | 501.96 | 316.17 | 18.01 |
| 1000 | 6500 | 42.46 | 4.99 | 3028.74 | 3645.43 | 21.50 | 1451.27 | 544.11 | 326.95 | 18.41 |
| 1000 | 7000 | 43.38 | 4.42 | 3211.73 | 3919.46 | 20.57 | 1423.18 | 521.60 | 329.16 | 17.40 |
| 1000 | 7500 | 45.91 | 4.48 | 3507.97 | 4291.63 | 23.89 | 1447.30 | 507.35 | 338.57 | 17.07 |
| 1000 | 8000 | 43.38 | 4.59 | 3656.90 | 4586.98 | 21.57 | 1417.73 | 511.91 | 364.30 | 18.13 |
| 1000 | 8500 | 46.58 | 05.03 | 3863.04 | 4916.73 | 21.40 | 1492.24 | 512.49 | 437.60 | 18.40 |
| 1000 | 9000 | 47.30 | 4.43 | 4042.96 | 5237.41 | 21.21 | 1424.07 | 526.02 | 497.90 | 18.12 |
| 1000 | 9500 | 46.68 | 4.43 | 4605.68 | 5490.45 | 20.37 | 1463.82 | 526.39 | 436.37 | 19.46 |
| 1000 | 10000 | 45.80 | 4.20 | 4889.18 | 5877.01 | 20.03 | 1485.12 | 534.18 | 464.18 | 17.52 |

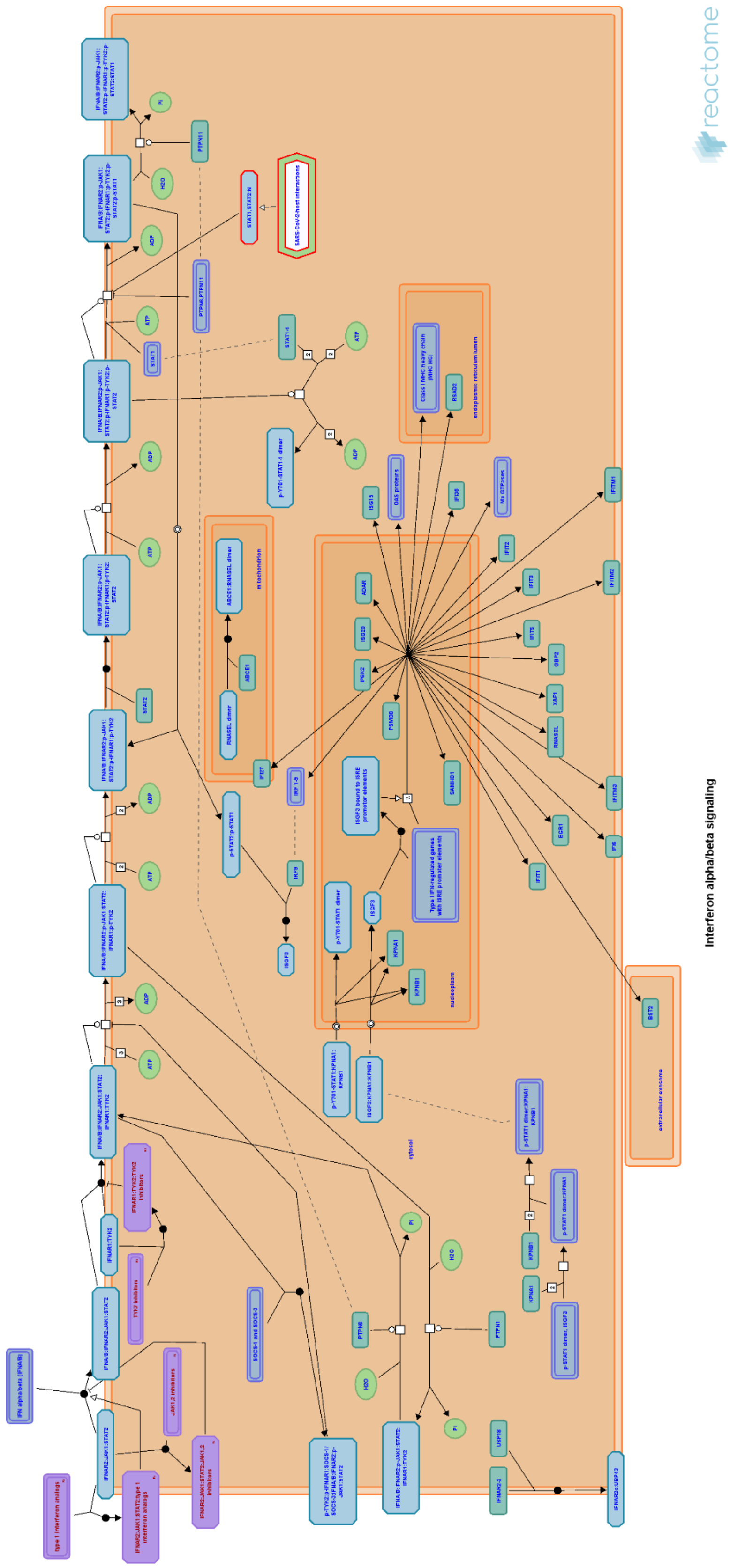


*Figure S1. Overview of the Reactome Interferon alpha/beta signaling pathway. Downloaded from Reactome Database with Reactome Viewer.*

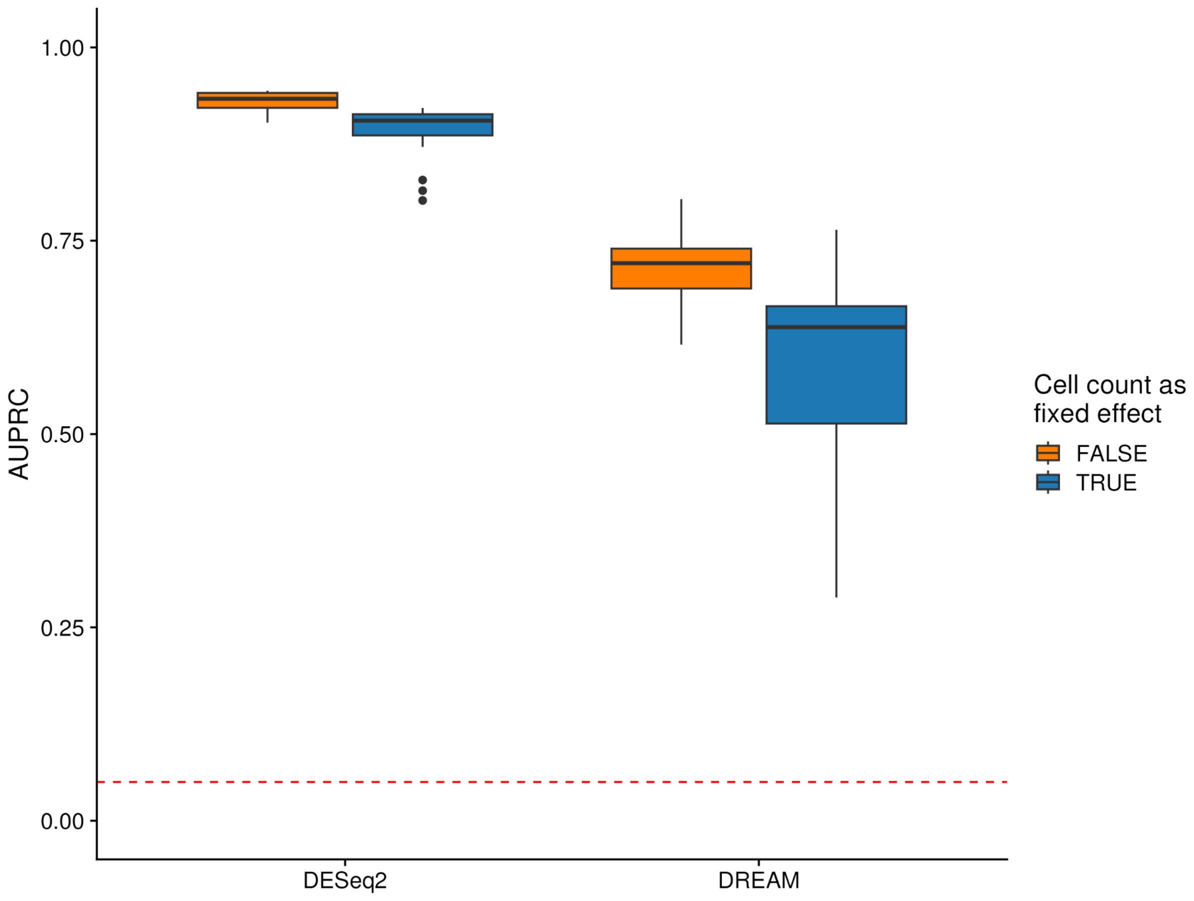


*Figure S2. Cell count per pseudobulk sample as fixed effect. The number of cells each pseudobulk consists of was incorporated as a fixed effect into the formula for the parametric pseudobulk methods DESeq2 and DREAM to check whether this increases the performance. The random baseline is indicated by the dashed line.*

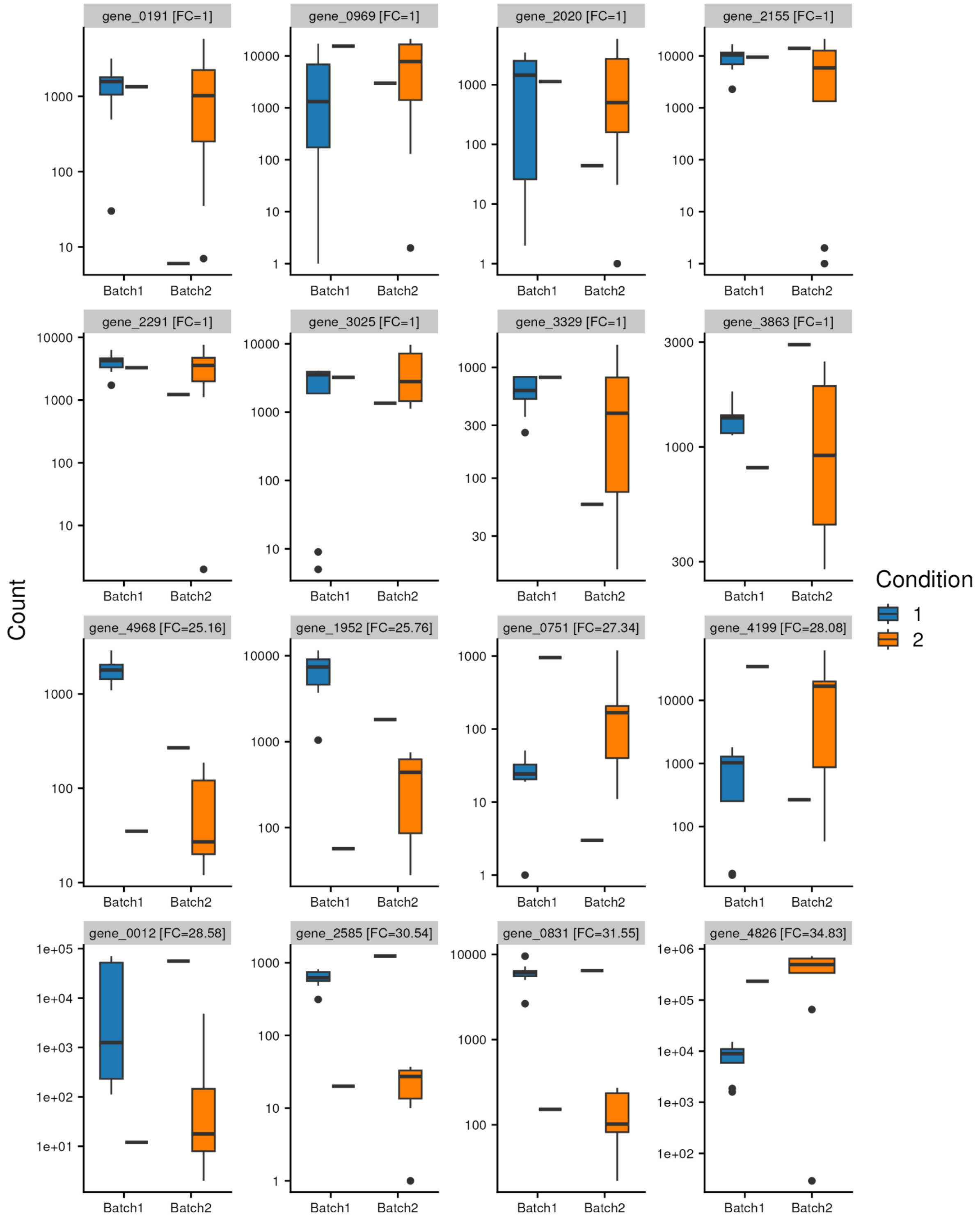


*Figure S3. Gene expression counts across batches and conditions in the unbalanced atlas setting. The figure shows the simulated expression per batch and condition for 8 randomly selected genes simulated not to be differentially expressed (log fold change = 0; top two rows) and the 8 genes with the highest differential expression by simulated log fold change (bottom two rows). The plot is based on the pseudobulked unbalanced atlas setting, each point refers to the gene count of one sample.*

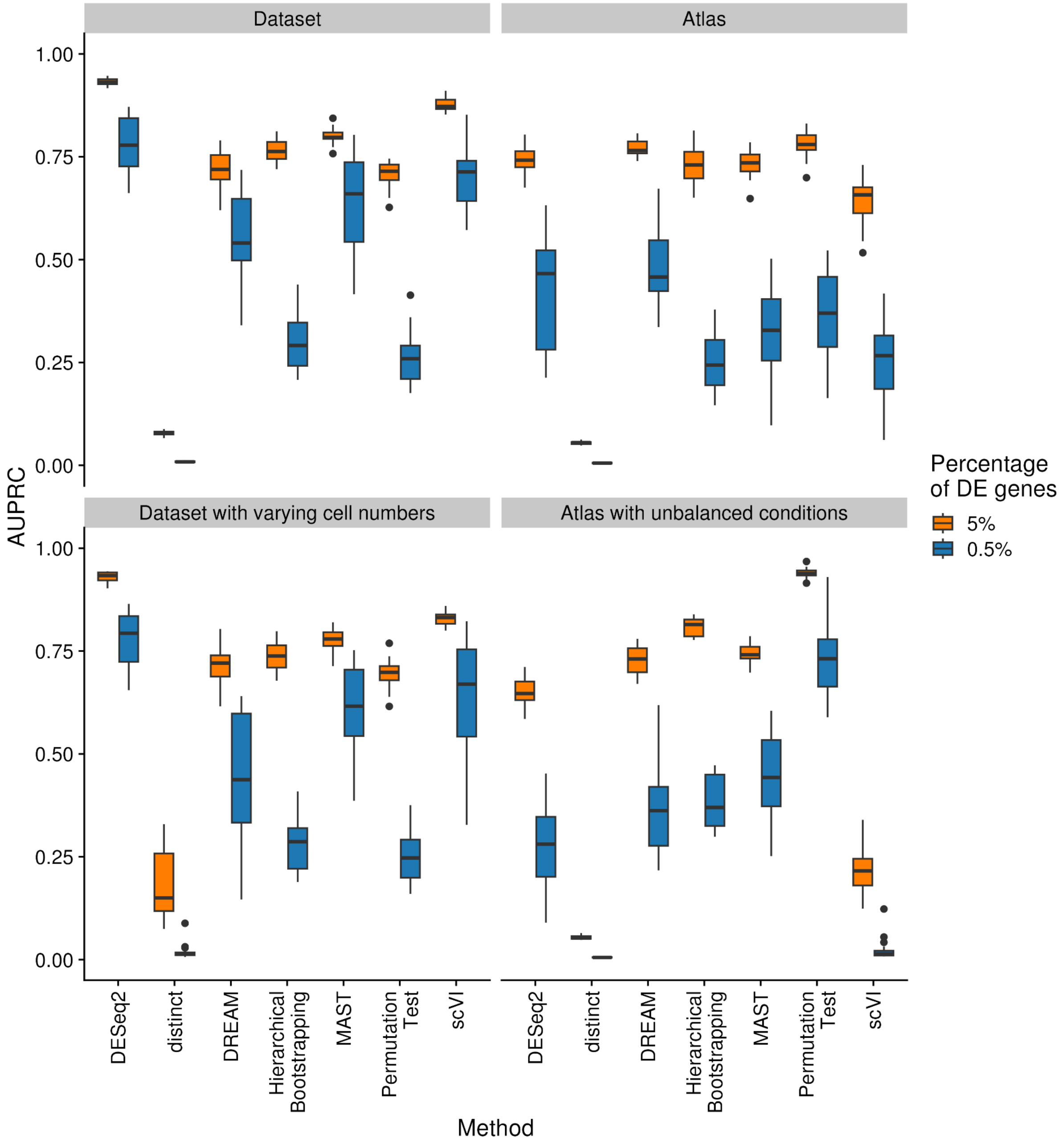


*Figure S4. Performance differences with varying DE gene percentage. Performance of all methods on all scenarios with different amounts (5% or 0.5%) of differential expressed genes. The scenarios refer to the four main simulation scenarios.*

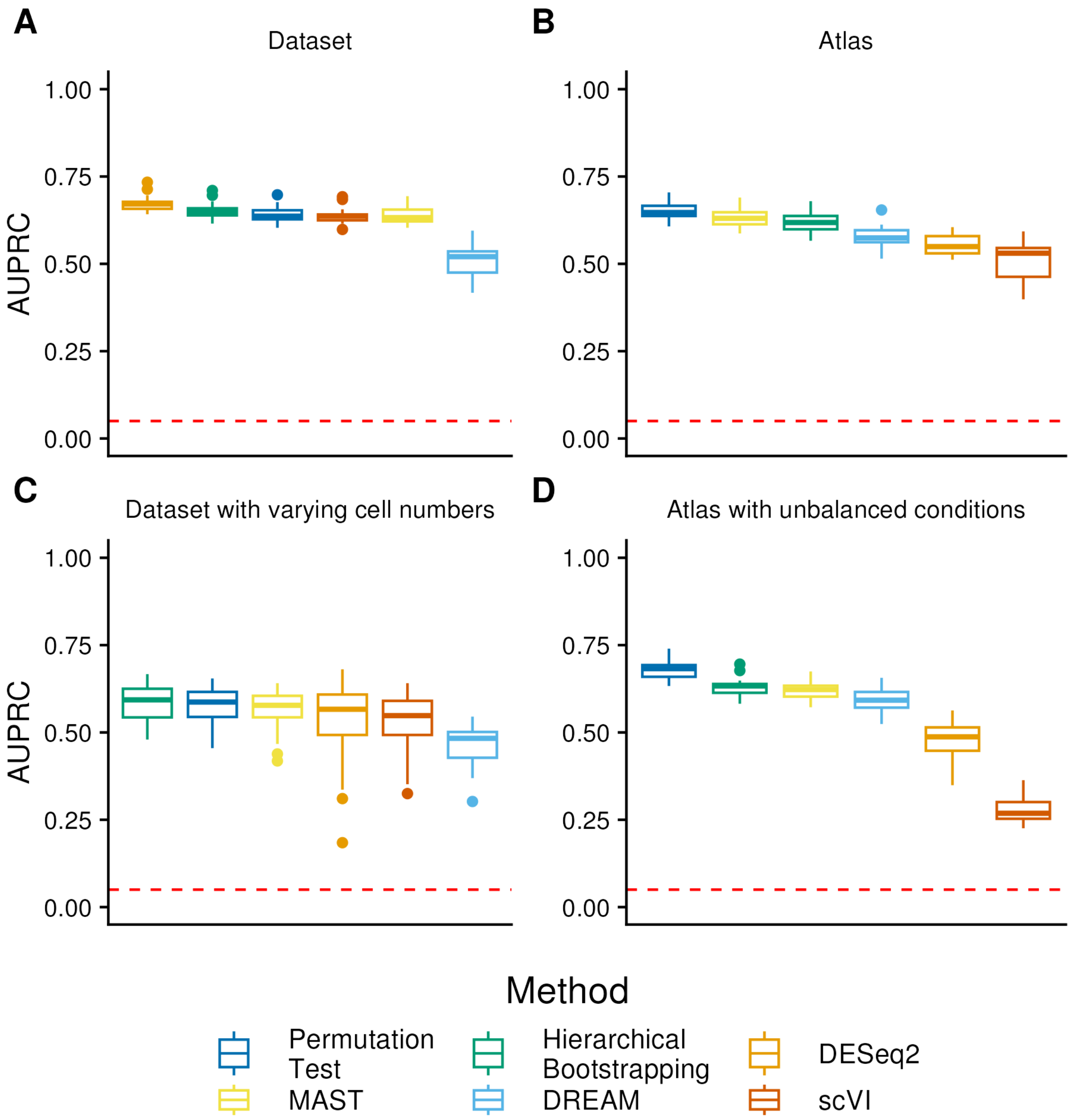


*Figure S5. Performance of the methods on the simulated scenarios after filtering for the top 10% of highly variable genes. Shown is the performance of the methods on the simulated scenarios over 20 independent simulations and runs each for (A) the Dataset scenario, (B) the Atlas scenario, (C) the dataset scenario with varying cell numbers per sample, and (D) the Atlas scenario with unbalanced conditions. Before the methods were applied to the simulated scenarios, the datasets were filtered for 10% of the highly variable genes. Removed differentially expressed genes are classified as false negatives. The methods were sorted based on their performance. The random baseline is indicated by the dashed line.*

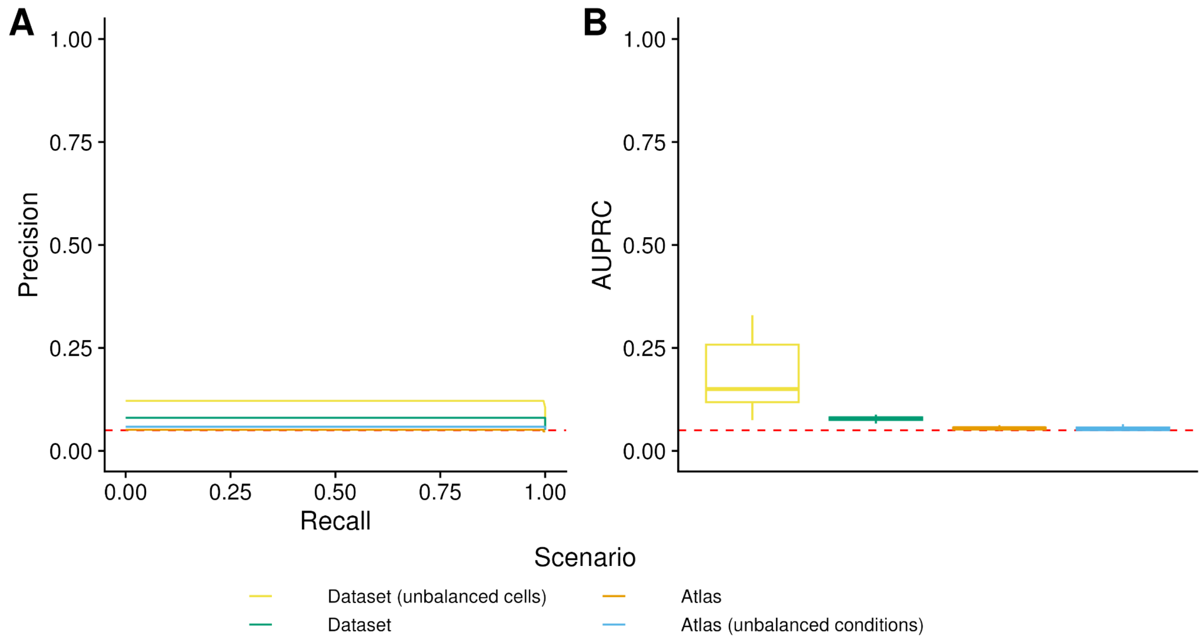


*Figure S6. Performance of distinct on the simulated scenarios. Distinct showed baseline performance in almost all simulated scenarios and was therefore excluded from the main benchmark. It assigns the minimal p-value to the majority of the genes and which makes them hard to rank.*

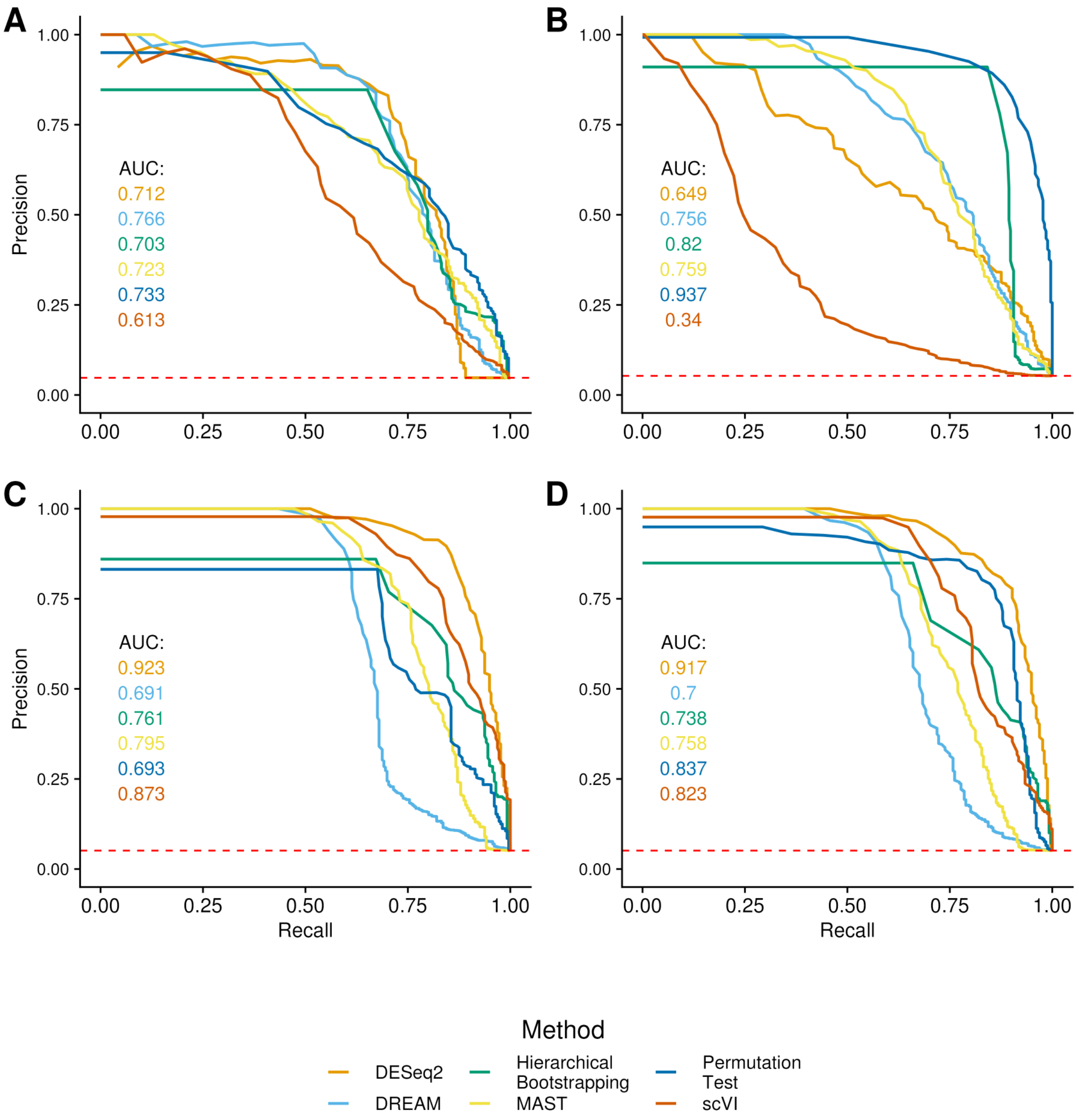


*Figure S7. Precision-recall plot for all scenarios on a single simulation. Shown are the precision-recall plots for one of the 20 simulations based on the predictions of all methods except distinct. (A) shows the data for the atlas scenario, (B) for the atlas scenario with unbalanced conditions, (C) for the dataset scenario and (D) for the dataset scenario with variable cell numbers.*

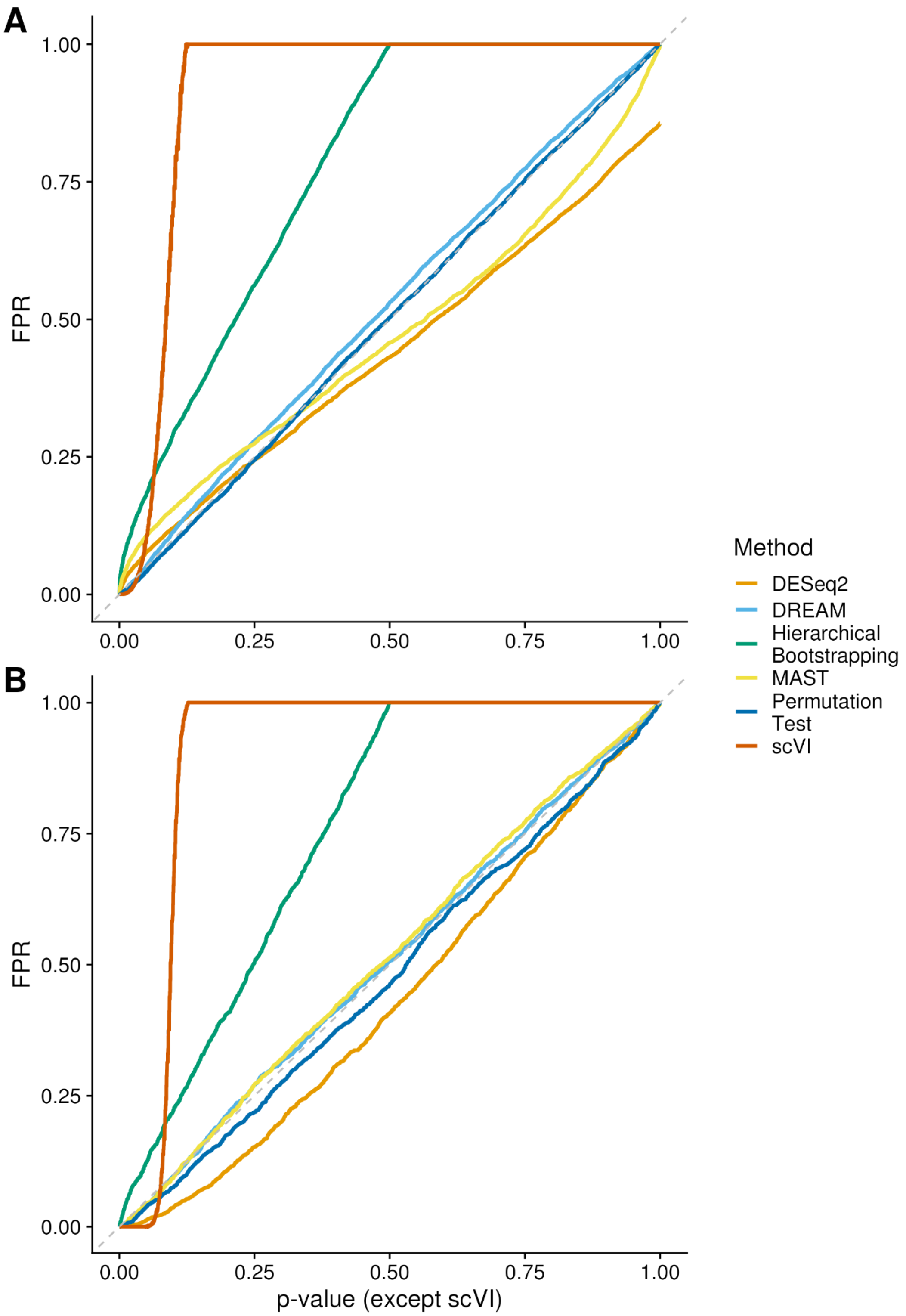


*Figure S8. Negative control based on raw p-values and false positive rate (FPR). Negative control based on (A) the simulated atlas scenario from Fig. 7 without zoom with no differential expression and (B) the real data scenario from the Seurat vignette. The real data has been permuted to eliminate differential expression across conditions. Since scVI does not generate traditional p-values but instead uses a Bayesian decision rule to determine differential expression, we assessed its classification results across 2000 equidistant cutoffs between 0 and 1, calculating false positives from these outcomes.*

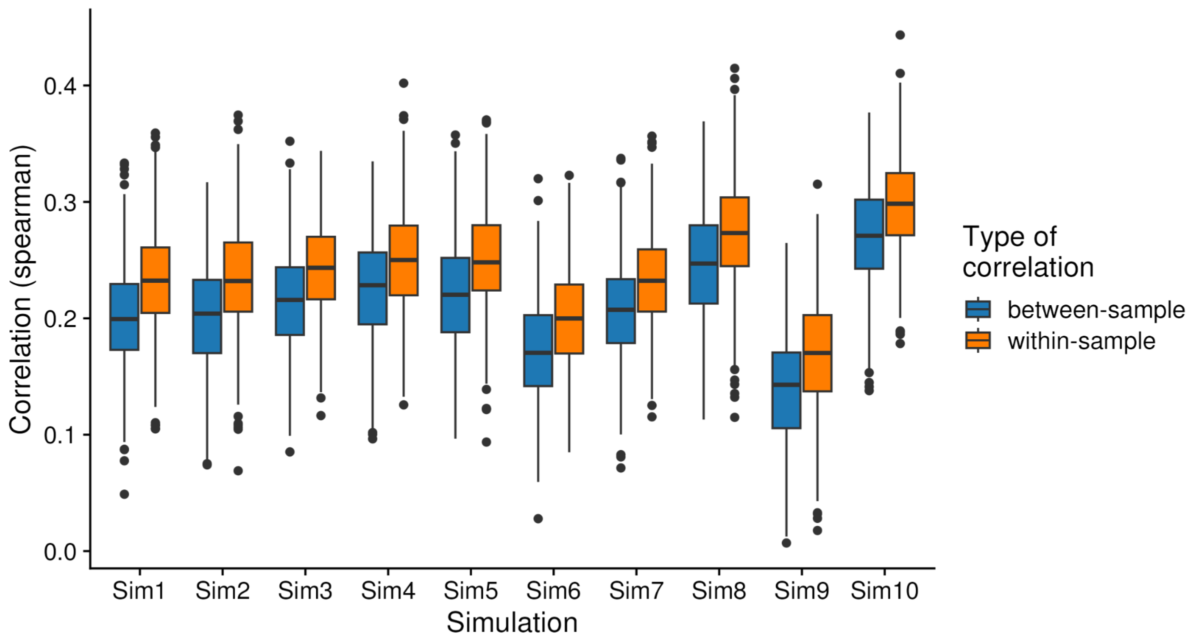


*Figure S9. Within- and between-sample Spearman correlations on the dataset scenario.The correlations are calculated based on the expression of two cells and depict the similarity of the expression of these cells to each other. The median of the correlations within a sample is always larger than the median across samples and therefore indicates a higher similarity of cells within a sample. The middle line represents the median, and the lower and upper limits of the boxes specify the 25% quartile and the 75% quartile. The whiskers extend to the largest/smallest correlation point, within 1.5 × interquartile range. Method adapted from Ref. [1]*
